# Newly identified COPD GWAS protein interactors reveal a potential disease network module

**DOI:** 10.64898/2026.09.17.752466

**Authors:** Dávid Deritei, Hiroyuki Inuzuka, Chengyue Zhang, Peter J. Castaldi, John M. Asara, Shariq Madha-Krause, Robert L. Moritz, Michael H. Cho, Kimberly Glass, Wenyi Wei, Edwin K. Silverman

## Abstract

**Background:** Chronic obstructive pulmonary disease (COPD) is a complex, heterogeneous disease influenced by genetic and environmental factors. However, the interactions between COPD risk genes and their collective role in COPD susceptibility remain largely elusive. We hypothesize that the protein-protein interaction (PPI) network of genes associated with COPD risk can provide molecular insight into the mechanisms of COPD pathogenesis.

**Methods:** We used affinity purification mass spectrometry (AP-MS) to detect protein-protein interactions of six well-established COPD GWAS gene products (AGER, FAM13A, FBXO38, HHIP, IREB2, MFAP2) in two relevant lung cell lines (IMR90, 16HBE). These genes are located in significant COPD GWAS regions, and their physiological role in COPD has been previously confirmed by functional studies. We analyzed the impact of the newly identified interactions in contextual, cell type-specific PPI networks built from the combination of publicly available PPI data (HUBRIS) and cell type-specific gene expression data.

**Results:** Using AP-MS, we identified 482 unique interactors across 701 newly identified protein interactions with the six GWAS gene products in at least one of the cell lines (AGER: 122, FAM13A: 83, FBXO38: 71, HHIP: 108, IREB2: 8, MFAP2: 309). Subsequent network analysis incorporating these new PPIs with public PPI databases revealed: (1) the new interactions significantly reduce the network distance between known COPD GWAS gene products; (2) 5.2% of the newly identified interactors are differentially abundant between COPD cases and controls in lung proteomic data from the Lung Tissue Research Consortium (LTRC); and (3) 41 (14.3%) of the newly identified interactors were directly connected to more than one of the six COPD GWAS genes. We identified a tightly connected core network of COPD GWAS gene products and protein biomarkers with intersecting signals; this network consists of 8 proteins, including 5 of the 6 GWAS gene products (AGER, FAM13A, HHIP, FBXO38, and MFAP2) as well as CAVIN1, TGM2, and HSPA1A.

**Conclusions:** By combining AP-MS experimental data, multiple types of “omics” data, and network analysis, we have constructed a disease network module for COPD. This module can be a foundation for understanding the collective influence of COPD GWAS gene products in disease pathogenesis.

## Introduction

Chronic obstructive pulmonary disease (COPD) is one of the leading causes of morbidity and mortality worldwide and represents a major global public health concern [1,2]. COPD is a complex disease arising from the interaction between genetic susceptibility and environmental exposures, notably cigarette smoke and biomass fuel exposure. Clinically, the disease is defined by persistent airflow limitation identified by spirometry, yet pathologically it manifests as a heterogeneous spectrum of phenotypes including airway remodeling, chronic inflammation, and alveolar destruction (emphysema). This heterogeneity suggests that multiple molecular mechanisms contribute to disease development and progression [3,4]. Genome-wide association studies (GWAS) have identified more than 80 loci associated with COPD susceptibility and many others associated with related traits such as lung function and emphysema [5–8]. The combination of genetic association and functional studies has highlighted multiple genes likely to be involved in COPD susceptibility, including *HHIP* [9]*, FAM13A* [10]*, IREB2* [11]*, AGER* [12]*, MFAP2* [13]*, FBXO38* [14]*, DSP* [15]*, FBLN5* [16]*, NPNT* [16]*, SFTPD* [17]*, TET2* [18]*, TGFB2* [19]*, MMP12* [20], and *MMP1* [21]. Functional follow-up studies have begun to elucidate the biological roles of individual COPD GWAS genes in processes such as response to smoke exposure [22], immune response [23], and iron metabolism [11]. However, despite these advances, the *collective* role of multiple COPD risk genes in influencing disease susceptibility remains poorly understood.

A growing body of evidence suggests that genes associated with complex diseases do not act in isolation, but instead likely form interconnected molecular networks. In such networks, disease-associated genes tend to cluster in close network proximity, with their protein products interacting directly or converging on shared pathways and biological processes [24,25]. Protein-protein interaction (PPI) network-based approaches have successfully revealed cell type- and disease-specific interactomes in neurodevelopmental disorders such as Autism spectrum disorders (ASDs) [26] and schizophrenia [27]. Moreover, bioinformatic studies of COPD GWAS loci similarly suggest functional relationships among risk genes, yet the physical and cell type-specific nature of these interactions has remained largely unexplored [28].

Protein–protein interaction networks offer a framework for bridging the gap between genetic association and biological mechanisms. However, most available PPI resources lack cell type specificity and do not involve lung-derived cells. To address this limitation, we previously developed HUBRIS, a network combining eight public PPI databases, and integrated it with cell type-specific transcriptomic data to contextualize experimental interaction data generated by our own group [29,30]. Specifically, we identified protein interactors of the protein produced from the COPD GWAS gene *HHIP* in two lung cell lines (IMR90 and 16HBE) and demonstrated that newly identified interactors shortened the network distance between HHIP and other COPD-associated gene products, revealing novel network paths enriched for biological processes relevant to COPD pathogenesis [29].

We hypothesized that systematic experimental mapping of PPIs for multiple well-established COPD GWAS gene products would reveal a shared disease sub-network underlying COPD susceptibility. Building on our previous work on *HHIP*, in the present study we used affinity purification mass spectrometry (AP-MS) to identify protein-protein interactions for products of five additional COPD GWAS genes--*AGER, FAM13A, FBXO38, IREB2,* and *MFAP2* --in two lung-relevant cell lines representing lung fibroblasts (IMR90) and bronchial epithelial cells (16HBE). By integrating these experimentally derived interactions with curated public PPI networks (HUBRIS [30]), lung transcriptomic and proteomic data from COPD cases and controls, and network topology analyses, we identified a compact, COPD-relevant interaction module that has the potential to enhance our understanding of the collective, mechanistic behavior of COPD genetic risk factors in COPD pathogenesis.

## Results

### 1. Study Overview: Cell type- and COPD-specific PPI networks integrating multiple layers of omic data

To construct COPD-relevant PPI networks, we integrated newly identified PPIs from six COPD GWAS gene products (see Results 2-3) with multiple data sources using an expanded pipeline from [29] (Figure 1). As previously described in [29], public PPI data from eight curated databases were compiled into the HUBRIS network, providing a reference of known interactions. Cell line transcriptomic data were used to generate cell type-specific networks and analyze the impact of the newly identified edges on network topology (Results 4). Finally, we leveraged bulk lung proteomic data (Madha-Krause, manuscript in preparation) from the Lung Tissue Research Consortium [31] to identify proteins correlating with the GWAS gene products or differentially abundant in COPD versus control lungs (Results 5-7). These datasets were combined iteratively, producing a series of contextualized, cell type-specific PPI networks that integrate both experimental and publicly available information, thereby enabling downstream analyses of network distance, multiple GWAS bridging proteins, and COPD-relevant disease modules.

**Figure 1.**
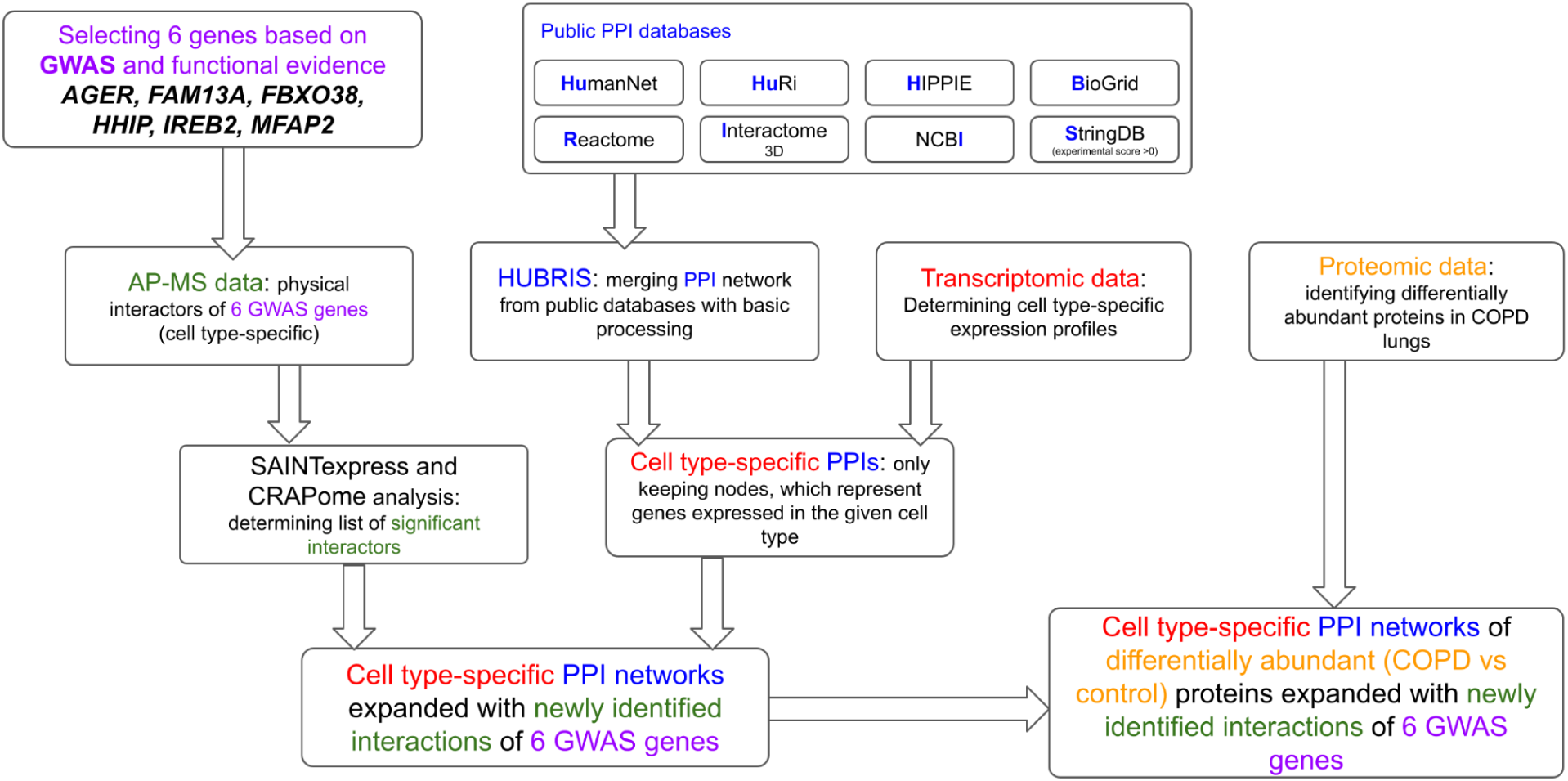
Network generation pipeline. We combined public data and data generated by our own research group to generate cell type- and COPD-specific protein-protein interaction (PPI) networks. The rectangles represent different stages of data processing, and arrows represent the combination of the different data types. The different colors highlight the different types of data or information. Purple: GWAS data and literature knowledge that informed the selection of the 6 COPD GWAS genes; Blue: public PPI data from 8 databases compiled into HUBRIS; Green: lung cell line specific (IMR90 & 16HBE) AP-MS data of newly identified interactors of the 6 GWAS gene products; Red: Transcriptomic (RNA-Seq) data from IMR90 and 16HBE cell lines; Orange: proteomic data generated from the Lung Tissue Research Consortium (LTRC) identifying differentially abundant proteins between COPD and control lungs (bulk tissue)

### 2. Affinity purification mass spectrometry identifies protein interaction partners of six COPD GWAS gene products in lung-relevant cell types

To systematically characterize protein-protein interactions (PPIs) of COPD risk genes, we performed affinity purification mass spectrometry (AP-MS) for five well-established COPD GWAS gene products: AGER, FAM13A, FBXO38, IREB2, and MFAP2; these results were combined with our previous HHIP analyses [29]. To ensure consistency with our prior work, we utilized the same experimental framework previously used to map HHIP protein interactions in lung-relevant cell types [29]. Specifically, we used 16HBE human bronchial epithelial cells, immortalized with SV40 large T antigen, and IMR90 fibroblasts derived from human fetal lung tissue and immortalized via hTERT expression. These cell lines represent epithelial and mesenchymal compartments, respectively, that are both critically involved in COPD pathogenesis.

Across all six COPD GWAS gene products and both lung cell lines, we identified a total of 701 protein-protein interactions involving 482 unique proteins using the Significance Analysis of INTeractome (SAINTexpress) validation tool [32]. To exclude noisy and non-specific interactions, we applied an additional layer of filtering based on the Contaminant Repository for Affinity Purification (CRAPome [33]) database (see Methods). After inclusion of the expanded CRAPome control set, 337 high-confidence interactions involving 286 unique proteins remained significant (Table 1). Most subsequent analyses are conducted with both sets: all interactors identified by SAINTexpress analysis and a more conservative set resulting from additional CRAPome filtering. In the results reported in this paper we prioritize the CRAPome filtered set, unless otherwise stated. Most of these interactions represent previously unreported PPIs, as only 3.7% (7.7% with SAINTexpress only) of the 337 (701 with SAINTexpress only) interactions are also present in the HUBRIS network. Only 5.9% (12.2% SAINTexpress only) of these interactions were observed in both lung cell lines.

**Table 1.** Number of experimentally identified protein-protein interactors of the six COPD GWAS genes. The rows of the table separate the number of cell type-specific interactions for each GWAS gene product. The first number in each cell represents the number of interactions after SAINTexpress validation with subsequent CRAPome filtering, and the second number in parentheses is the number of significant interactions *without* additional CRAPome controls (SAINTexpress only).

| Cell line/GWAS gene | MFAP2 | FBXO38 | AGER | IREB2 | FAM13A | HHIP | Total (columns) |
| --- | --- | --- | --- | --- | --- | --- | --- |
| IMR90 only | 14 (49) | 17 (36) | 78 (104) | 1 (1) | 26 (79) | 8 (74) | 144 (343) |
| 16HBE only | 157 (208) | 4 (20) | 1 (6) | 3 (7) | 0 (0) | 8 (31) | 173 (272) |
| both cell lines (intersection) | 9 (52) | 7 (15) | 3 (12) | 0 (0) | 0 (4) | 1 (3) | 20 (86) |
| Total (union) | 180 (309) | 28 (71) | 82 (122) | 4 (8) | 26 (83) | 17 (108) | 337 (701) |
| Overlap with HUBRIS (any cell line) | 0 (0) | 9 (10) | 3 (3) | 3 (3) | 10 (10) | 0 (0) | 25 (26) |
| Percentage of overlap (intersection with HUBRIS %) | 0% (0%) | 35% (14%) | 3.7% (2.5%) | 75% (38%) | 38% (12%) | 0 (0) | 7.7% (3.7%) |

The distribution of identified interactions differed substantially across COPD GWAS bait proteins and cell types (Figure 2 and Table 1). Several COPD GWAS gene products exhibited cell type-specific interaction profiles. In particular, AGER and FAM13A showed a greater number of IMR90-specific interactions, while MFAP2 displayed an enrichment of interactions specific to 16HBE cells. These patterns suggest that the interaction networks of COPD GWAS genes are likely shaped by cellular context, consistent with the heterogeneous tissue involvement observed in COPD. Moreover, the number of newly identified interactors did not reflect the data available in public PPI resources. Comparison with the HUBRIS network revealed marked variability in existing PPI coverage across the six GWAS gene products. For example, FBXO38 and IREB2 yielded relatively few interactors in the AP-MS experiments, but they are connected to more than 50 and 100 interaction partners in HUBRIS, respectively. For these two proteins, the overlap of the newly identified interactions with the public data is also higher (Table 1), indicating lower lung-specificity. It is worth noting that in most public PPI databases, the cellular context of the interactions is not documented. The full list of interactors with detailed meta-data is available in Supplementary Table 1.

**Figure 2:**
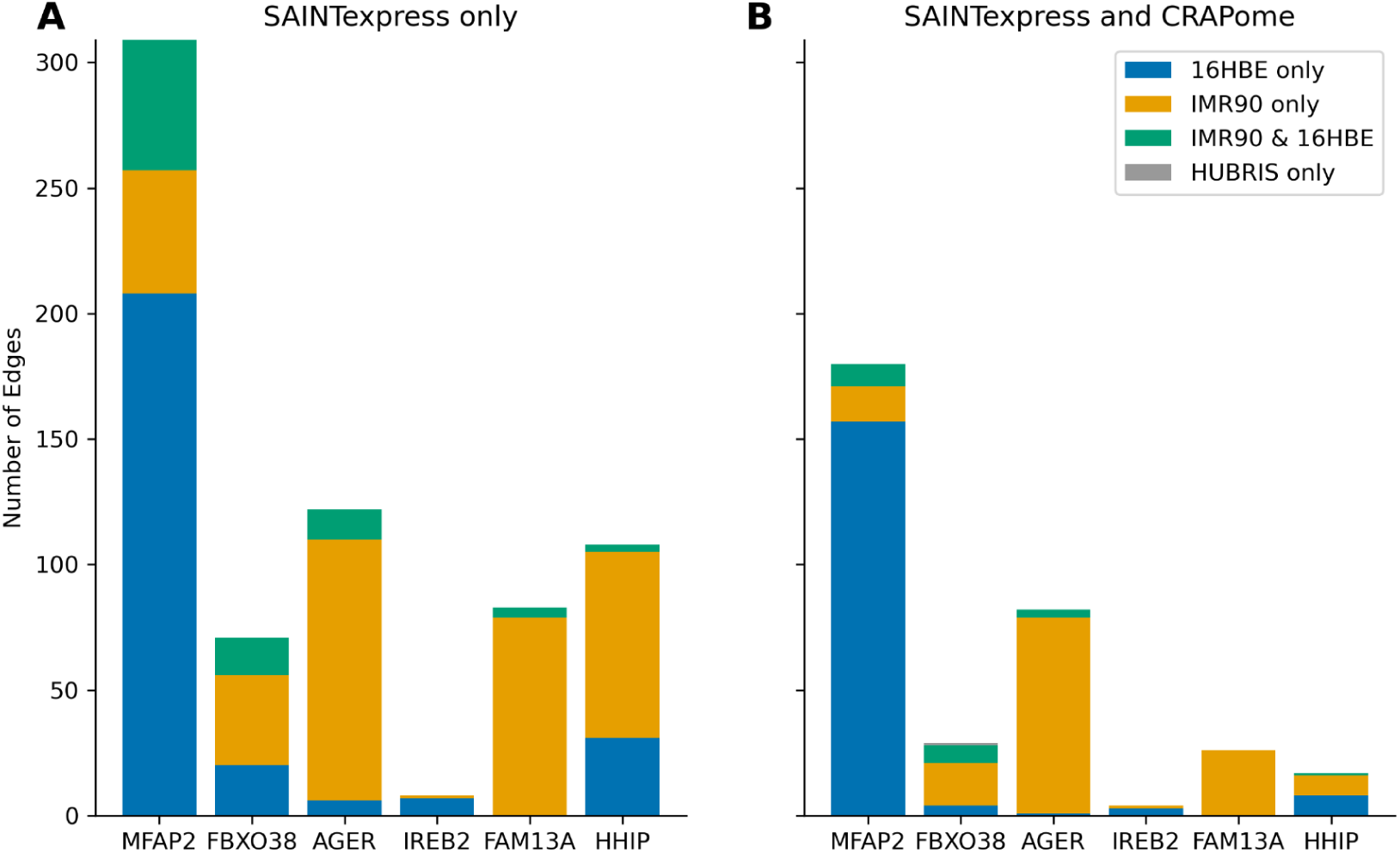
Distribution of newly identified cell type-specific and publicly available interaction counts for the six COPD GWAS gene products. Colors represent the origin of the interactions (cell type or public database) Left: number of interactions identified with the SAINTexpress pipeline without CRAPome controls. Right: number of interactions identified with additional CRAPome-based filtering.

### 3. Some newly identified interactors are implicated as likely COPD GWAS gene candidates

Of the genes encoding the 482 newly identified interactor proteins, 33 (6.85%) have their transcription start site (TSS) within +/-1 megabase pair (Mb) of a lead single nucleotide polymorphism (SNP) from 82 COPD GWAS loci, identified by [34] (Supplementary Table 2). Of these 33, most (28) are not the nearest protein-coding gene to the GWAS SNP; however, they may have functional relevance. The remaining 5 of the 33 genes are the nearest protein-coding genes to a COPD GWAS SNP; this includes HSPA4, which interacted with both AGER and MFAP2 (Table 2). Of the remaining 28, 4 genes (*HLA-B*, *HLA-C, HSPA1A* and *HSPA1L*) are within 1 Mb of the same SNP: rs2070600, the COPD GWAS SNP located in *AGER*. The gene products of *HLA-B*, *HLA-C* and *HSPA1A* also interact with *AGER*’s gene product in our new data. Moreover, *HSPA1A*, a gene encoding a heat shock protein that has a central role in our network analysis results (Results 7).

**Table 2:** Newly identified interactors that are potential COPD GWAS genes. The rows represent newly identified interactors between the proteins encoded by the genes on the left-most column (Newly identified interactor) and the right-most column (COPD GWAS gene product). The right-most column represents one of the 6 GWAS genes used as bait, and the left-most column represents genes that encode proteins that were identified as protein-protein interactors *and* are the nearest gene to a COPD GWAS SNP (second column). The rows highlighted in bold are interactions also validated after CRAPome filtering.

| Newly identified interactor | COPD GWAS SNP | TSS distance to SNP (base pairs) | Interaction identified in (cell line) | COPD GWAS gene product (protein) |
| --- | --- | --- | --- | --- |
| **TNPO1** | **rs34651** | **31866** | **IMR90** | **AGER** |
| HSPA4 | rs62375246 | 51356 | both | AGER |
| **HSPA4** | **rs62375246** | **51356** | **16HBE** | **MFAP2** |
| **PRSS23** | **rs117261012** | **57340** | **16HBE** | **MFAP2** |
| RPL23 | rs34727469 | 169039 | 16HBE | MFAP2 |
| **DDX1** | **rs10929386** | **174877** | **16HBE** | **MFAP2** |

### 4. Newly identified interactions reduce network distance between COPD GWAS gene products

To assess whether the newly identified protein-protein interactions alter the network relationships between COPD GWAS genes, we examined their effect on the pairwise network distances among the six GWAS gene products within the cell type-contextualized PPI networks. Incorporation of the experimentally derived interactions remaining after CRAPome filtering resulted in a significant reduction in network distance (length of the shortest path) between the GWAS gene products compared to a null model in both cell type-specific networks (p = 0.038 for 16HBE and p = 0.004 for IMR90). However, significant differences were not observed when all interactions (without CRAPome filtering) were included (Figure 3). The null distribution was generated using 1,000 random degree-preserving rewirings in which the non-GWAS interactor node of each newly added interaction was randomly reassigned to a node of similar degree, preserving the degree of both the GWAS genes and the randomly assigned neighbors. This indicates that the more conservative CRAPome filtered edge subset may have higher biological relevance. Therefore, in all subsequent analyses, unless otherwise noted, we use the CRAPome-filtered interaction set.

**Figure 3.**
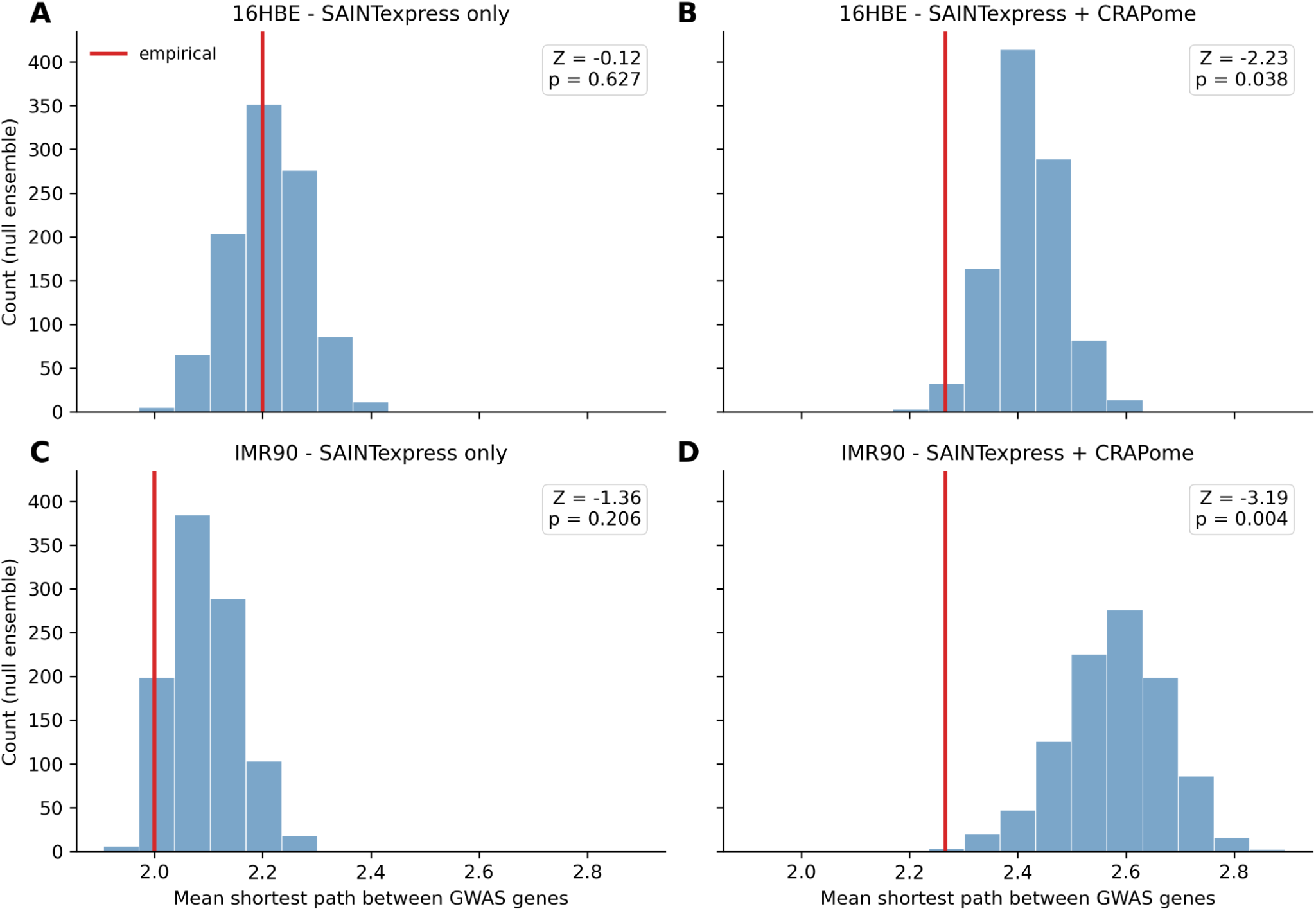
Experimentally identified edges reduce the network distance between COPD GWAS gene products after CRAPome filtering. For each combination of cell line (16HBE, IMR90) and edge-filtering criterion (SAINTexpress only, SAINTexpress + CRAPome), the mean shortest-path length between all pairs of the six COPD GWAS genes (MFAP2, FBXO38, AGER, IREB2, FAM13A, HHIP) was computed on the empirical network (red line) and compared to a null distribution (blue) of 1,000 degree-preserving randomizations. In each randomization, the non-GWAS endpoint of every new experimental edge was rewired to a randomly chosen node drawn from the same degree bin (deciles of the HUBRIS network degree distribution), holding the baseline network and the number of added edges fixed. The empirical distances fall below the null distribution for CRAPome-filtered edges in both cell lines (panels B, D), indicating that these edges bring the GWAS genes closer together than expected by chance, whereas edges derived using SAINTexpress only (without CRAPome filtering) show no such effect (panels A, C).

This statistically significant reduction in network distance demonstrates that the newly identified experimental PPIs are not isolated pairwise interactions. Furthermore, the mean shortest path distance after including the new interactions is ∼2.25, which means that the pairwise path-lengths between COPD GWAS proteins are mostly of length 2 (meaning one protein separating them; there are no direct interactions between the GWAS proteins, i.e., of path-length 1). This implies that the reduction in average path length is due in part to proteins that directly *connect to multiple COPD GWAS gene products, effectively acting as network bridges* (Figure 4). When taking CRAPome filtered interactions from either cell line, 41 (14.3%) of the newly identified interactors are directly connected to two or more of the six COPD GWAS genes. These multi-GWAS interactors create new paths (of two edges) between risk genes that were previously separated by multiple proteins in the network. The bridging proteins show cell-type specificity: the subnetwork of interactors that are bridging GWAS proteins *and* were identified in 16HBE cells (Figure 4A) is enriched for proteins involved in pathways including epithelial-mesenchymal transition (EMT) and complement (Figure 4C), while the IMR90 bridging network is enriched for TGB-β, AMPK, and HIF-1 signaling pathways (Figure 4B-C). Notably, both cell type-specific networks are enriched for the Hypoxia and Protein processing by the endoplasmic reticulum pathways, and share one protein (other than the GWAS proteins): THBS1. THBS1 was identified as a key gene/protein in multiple lung-function related GWAS studies (see Discussion).

**Figure 4.**
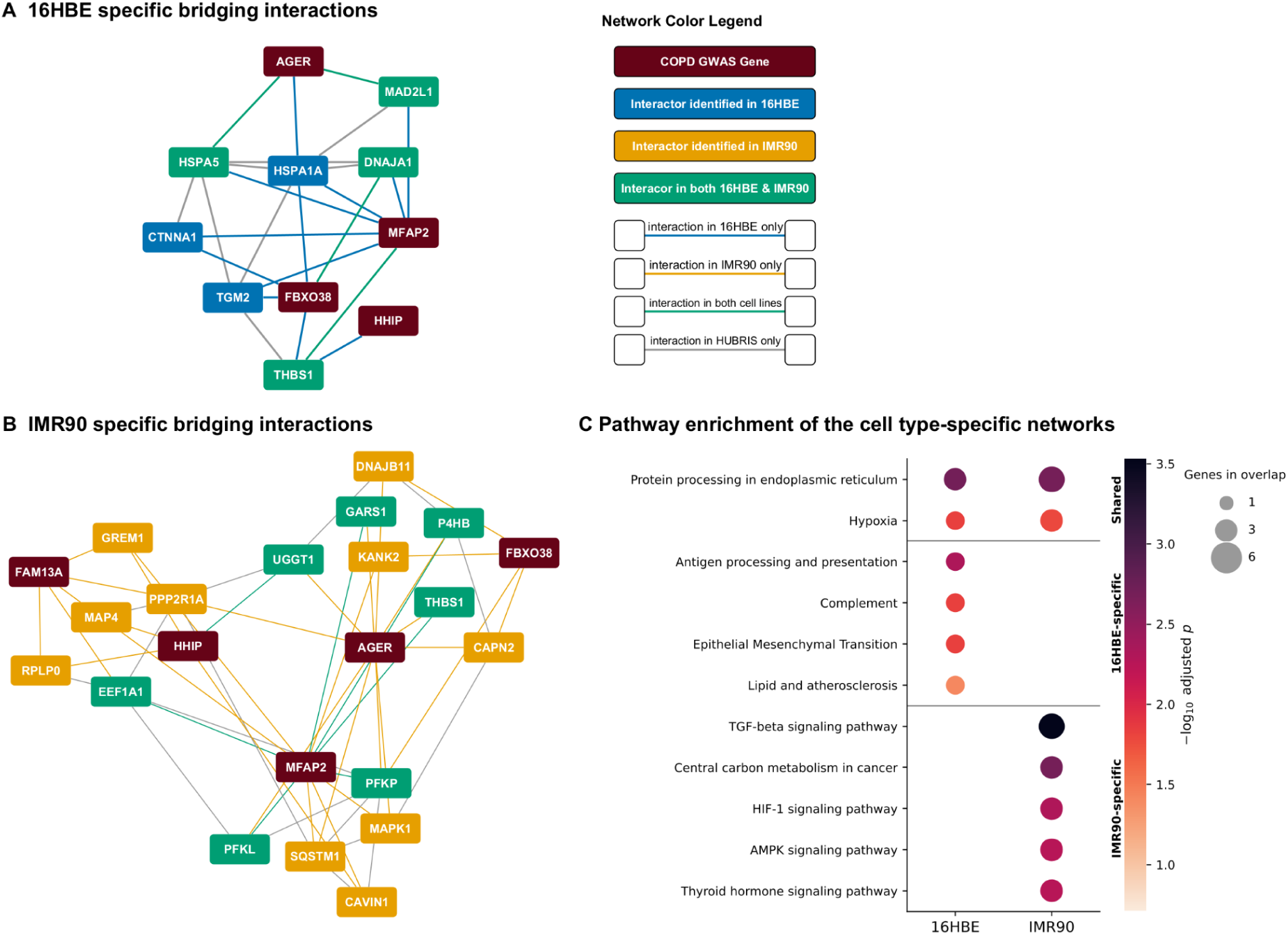
Cell type-specific networks of newly identified interactors that bridge multiple GWAS gene products. Nodes in the networks (both A and B panels) are newly identified interactors that connect to two or more COPD GWAS gene products (red nodes) in the same cell type. The colors of nodes represent the cell line in which they were identified as interactors, except for GWAS gene products (dark red). Layout is generated by a Force layout algorithm (yED 3.25.1) with slight manual editing. C-Functional enrichment of network nodes in Hallmark and KEGG pathways. The top 5 pathways per category are shown. The size of the dots is proportional to the overlap with the pathway(s), and the color is scaled with the -log10 of the adjusted p-value. Only terms with FDR < 0.05 are displayed. The COPD GWAS gene products were not included in the enrichment analysis.

### 5. Residual co-abundance of COPD GWAS proteins reveals biological context

We next investigated the COPD GWAS gene products and their protein interactors using data quantified from 4511 proteins, based on 452 COPD and 313 control lung tissue samples analyzed using mass spectrometry proteomics from the Lung Tissue Research Consortium (LTRC) (Madha-Krause, in preparation). Of the six COPD GWAS gene products in our study, only three, AGER, MFAP2, and FBXO38, were detected at the protein level in lung tissue, and all proteomic analyses are therefore restricted to these three GWAS gene products.

We adjusted the protein levels using the ordinary least squares method for age, sex, smoking status, smoking pack-years, ten ancestry principal components, and fifteen surrogate variables. The residual protein values were calculated by subtracting the fitted from the observed values. (for more details see Methods). Next, we calculated the residual Spearman correlation for each of the 3 COPD GWAS proteins with all other proteins. We found 1865, 1280, and 50 significantly (FDR < 0.05) correlated proteins with AGER, MFAP2 and FBXO38, respectively. We then performed pathway enrichment analysis for each GWAS protein by ranking significantly correlated (FDR<0.05) proteins by their residual correlation values with the GWAS protein (GSEA preranked) (Figure 5). For AGER, this analysis identified enrichment for cell-cell junction, plasma-membrane localization, caveola, and adherens junctions, consistent with AGER’s localization to the alveolar type I cell membrane. Strikingly, the 6 proteins with the highest correlation with AGER included the caveolar components EHD2 (FDR=3.94e-98, also a newly identified interactor of AGER in IMR90 cells), CAVIN2 (2.81e-79), CAV1 (1.23e-65) and CAVIN1 (FDR=6.34e-76, an experimentally identified interactor of both HHIP and MFAP2) (Supplemental Table 2). The MFAP2 analysis identified enrichment for extracellular matrix organization, collagen formation, elastic fiber, and microfibril terms, matching its role in elastic fiber assembly. In contrast, the proteins correlated with FBXO38 were enriched predominantly for lysosomal and antigen-presentation terms (lysosomal lumen, lytic vacuole, MHC class II antigen processing and presentation, and amino-acid catabolism). A small set of terms were shared across the analyses: proteins correlated with AGER and MFAP2 were jointly enriched for basement-membrane and myoblast-fusion/syncytium-formation terms, while proteins correlated with AGER and FBXO38 shared a lysosomal-lumen signature. These results indicate that the three GWAS proteins anchor largely distinct but partially overlapping biological modules in the lung proteome.

**Figure 5.**
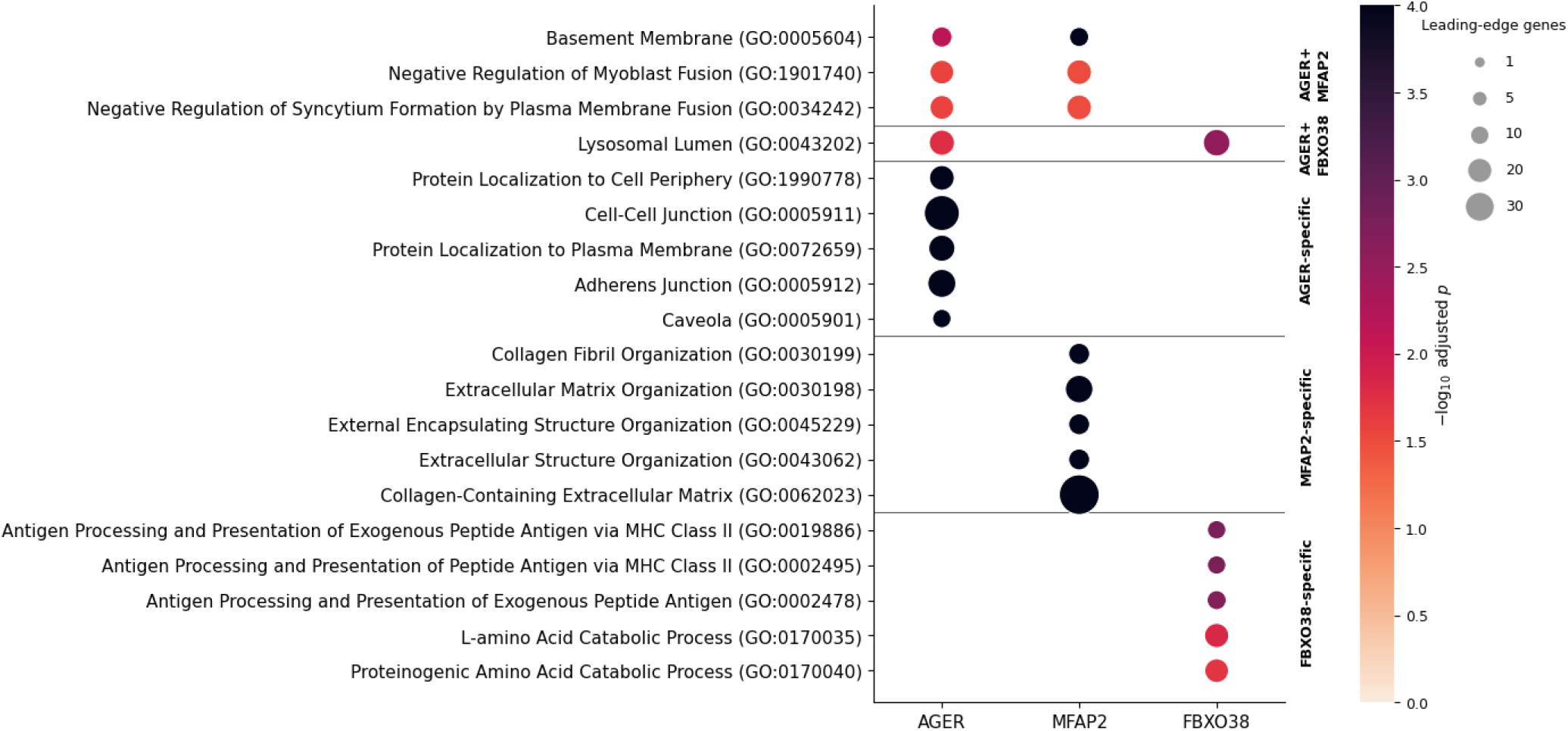
Pathway enrichment based on protein correlation reflects canonical functions. Gene-set enrichment analysis (GSEA preranked, 10,000 permutations) of each GWAS protein’s genome-wide residual correlations, ranked by signed Spearman correlation coefficient, against MSigDB Hallmark 2020, GO Cellular Component 2025, and GO Biological Process 2026. Enriched terms (FDR < 0.05) are grouped into blocks by the combination of GWAS proteins in which they are significant with the top five terms shown per block, ranked within each block by the minimum −log₁₀(FDR) across the contributing GWAS proteins. Dot color indicates −log₁₀(FDR-adjusted p), capped at the permutation resolution limit (FDR floor = 1e-04), and dot size indicates the number of leading-edge genes. Only terms with FDR < 0.05 are displayed.

We next asked whether the experimentally identified protein-protein interactions are enriched among proteins correlated with COPD GWAS gene products. Of the 1865 proteins that show significant correlation with AGER, 30 are also physical interactors of AGER in our data. These interactions are predominantly identified in IMR90 cells (90%) and are enriched for endoplasmic reticulum and apoptosis-related terms. Of the 1280 proteins significantly correlated with MFAP2, 47 are direct interactors, predominantly in 16HBE cells (76.6%) and are enriched for the epithelial to mesenchymal transition (Supplementary Figure S1; Supplementary Table 3). Given these results, for each GWAS gene product, we tested whether its experimental interactors ranked higher among its correlates than expected by chance (one-sided Wilcoxon rank-sum). Two cell type-specific signals emerged: MFAP2 interactors in IMR90 (n = 21; median rank 1131 vs 2256 background; p = 0.018) and AGER interactors in 16HBE (n = 3; median rank 539 vs 2255; p = 0.011), the latter was strong in effect but based on only three measured interactors (HSPA1A, HSPA5, and SEC61A1) and therefore is underpowered. The other GWAS protein-cell line combinations showed no significant signal.

Differential correlation analysis (COPD vs. control; Fisher Z-test, FDR < 0.05) identified eleven proteins whose correlation with AGER changed significantly with disease status (Supplementary Table 4). Of these, only one was a physical interactor of any of the COPD GWAS proteins: transglutaminase 2 (TGM2), an ECM-crosslinking enzyme identified as an interactor of FBXO38 and MFAP2 (Figure 4A). Of note, TGM2 was also identified as an interactor of AGER in the SAINTexpress-only analysis (without CRAPome filtering). TGM2 correlated positively with AGER in controls (r = 0.22), but this relationship was lost and reversed in COPD (r = −0.12); the differential correlation thus has an r-delta of -0.3341 (FDR = 0.0074).

### 6. Disease-associated network components reveal an AGER-centered caveolar module

Madha-Krause et al. identified 248 proteins with significant differential abundance between COPD and control lungs in the LTRC lung cohort (in preparation). The caveolar proteins we found to have the most significant co-abundance with AGER (Results 5, Supplementary Table 2) are also amongst the most significant differentially abundant proteins (EHD2, CAVIN1, CAVIN3, EHD1, and CAV1) between COPD and control identified by Madha-Krause et al. We intersected these differentially abundant proteins with the cell type-specific experimental interactors of the six COPD GWAS genes, identifying 16 proteins from our network that are implicated in COPD. We found that the differentially abundant proteins were not over-represented among the experimental interactors of the six GWAS genes (Fisher’s exact p = 0.46), indicating that the two signals are statistically independent.

Of the 16 identified proteins, the caveolar proteins (CAVIN1, EHD1, EHD2) form a tight correlation cluster with AGER and each other; this cluster is anchored around the physical interactions between AGER and EHD2 (Figure 6). This AGER-centered caveolar block is anti-correlated with a block of proteins made up of FHL2, PLD3, and CTSB, the former two of which also physically interact with AGER. The caveolar block also shows slight anti-correlation with the correlation block formed around MFAP2 (with TGM2 and PSMC6).

**Figure 6.**
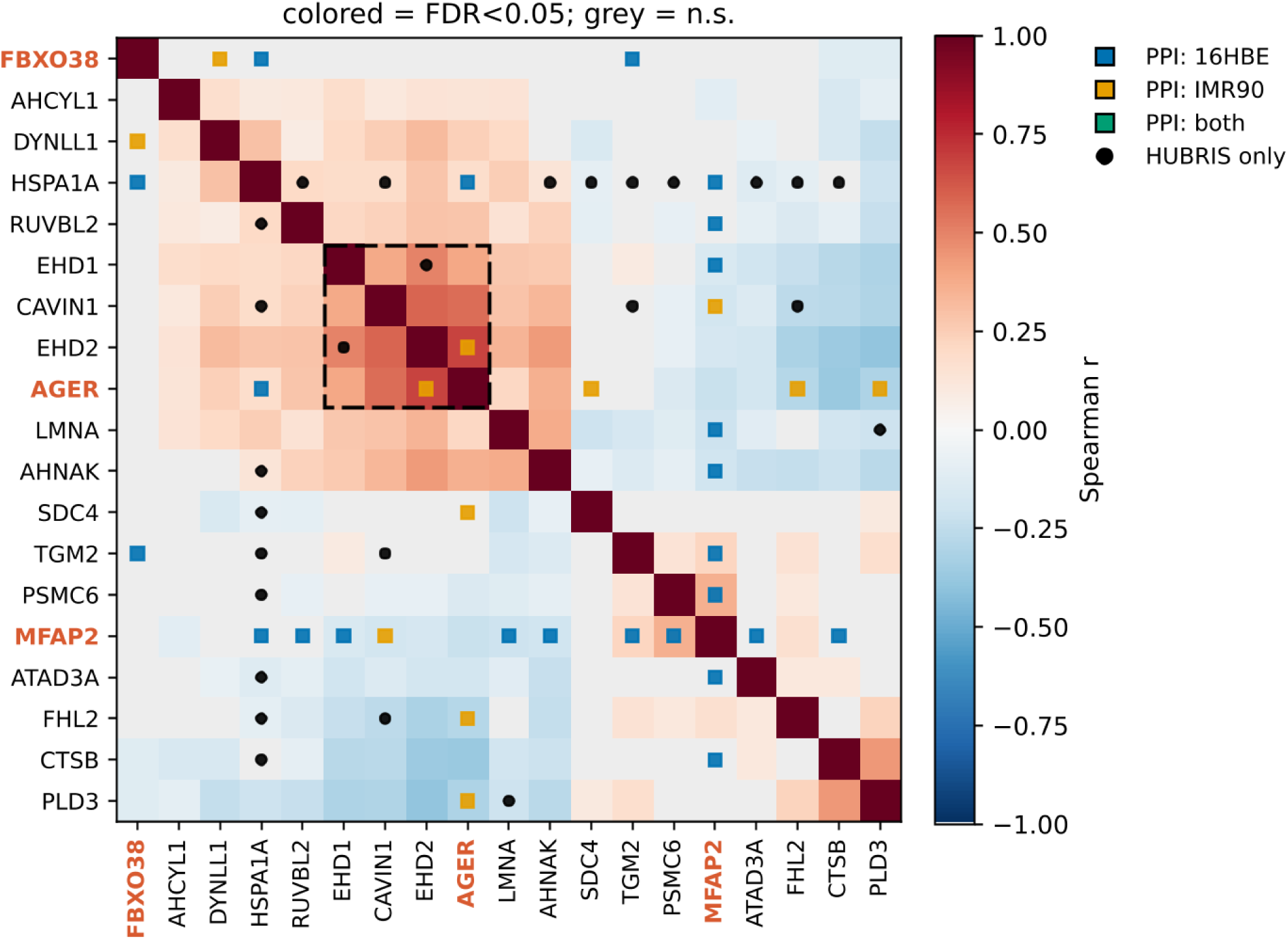
Residual protein correlation matrix of GWAS interactor proteins that are differentially abundant between COPD and control lungs. Spearman correlation matrix, computed on covariate-adjusted residuals across the COPD and control subjects in the lung tissue proteomic cohort (excluding IPF patients). Proteins were selected based on meeting two criteria: First, they are in the GWAS gene product network of interactors (16HBE, IMR90, CRAPome filtered, combined with HUBRIS, largest connected component); second, they are differentially abundant between COPD and control. Proteins are ordered by hierarchical clustering (average linkage on 1 − r), so co-expressed blocks appear contiguous along the diagonal. The dashed square highlights the caveolar block. Cells are colored by Spearman r only where the correlation is significant (FDR < 0.05); non-significant cells are shown in grey. GWAS gene products are labeled in orange. Colored squares mark experimentally identified physical interactions in our affinity-purification data; black circles mark interactions present only in the HUBRIS reference interactome.

### 7. Identification of a core COPD interaction network bridging GWAS gene products and lung proteomic changes

Finally, to further refine the set of disease-relevant interactions, we derived a core-network that intersects the following criteria: (1) only includes newly identified lung-specific PPIs of GWAS gene products (Results 2) that connect to two of more COPD GWAS gene products (Results 4, Figure 4); (2) only includes proteins that show differential protein abundance in COPD lung tissue (Results 6, Figure 6); and (3) forms a single connected component. The resulting core network is shown in Figure 7A.

**Figure 7.**
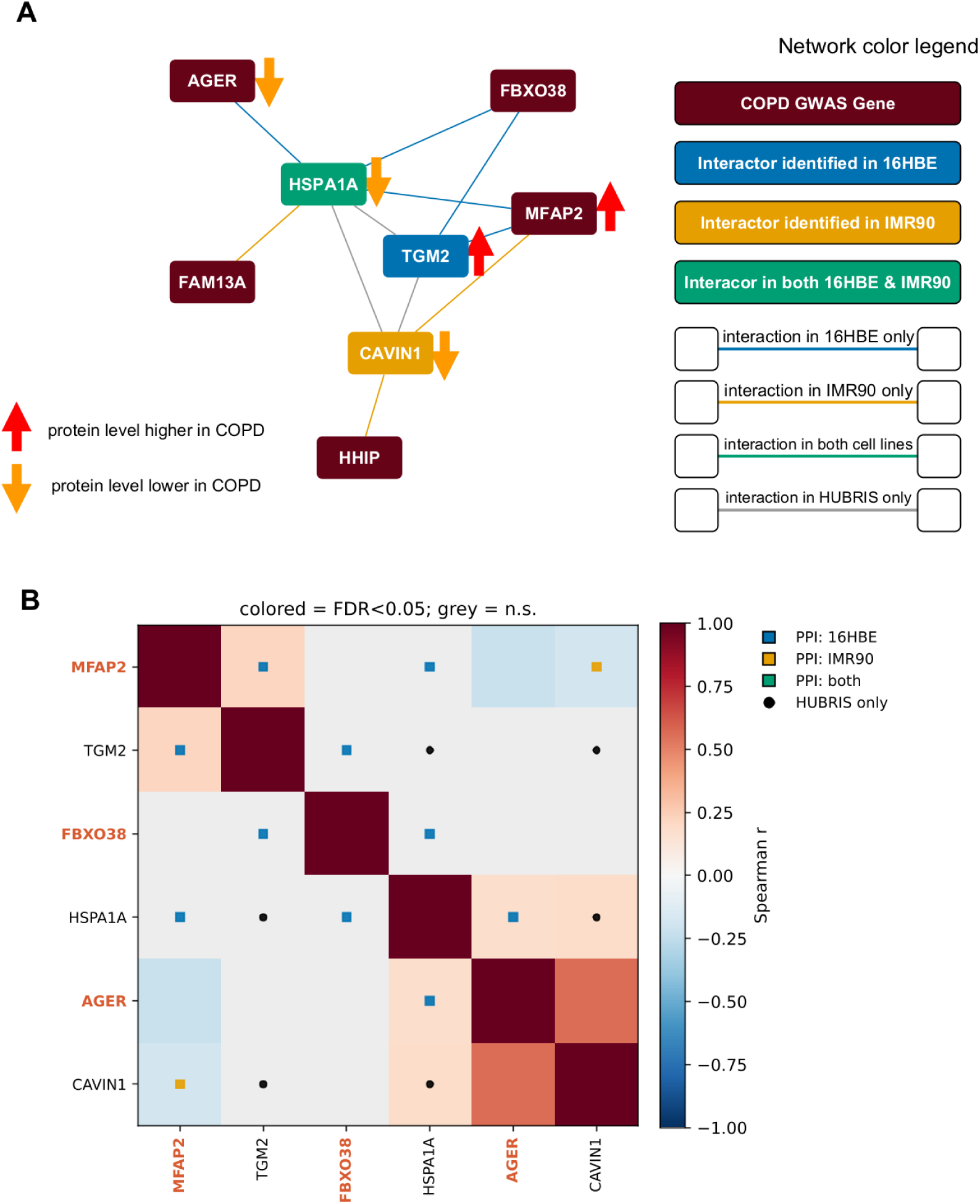
Core COPD protein-protein interaction network. (**A**) Every non-COPD GWAS node in the network is a newly identified interactor that connects to two or more COPD GWAS gene products (see Figure 4) and is a differentially abundant protein between COPD and controls in lung tissue. The colors of nodes represent the cell line they were identified in, except for GWAS genes (dark red). The arrow symbols beside the nodes represent whether the protein levels are significantly higher or lower in COPD compared to controls. Two of the GWAS gene products are also differentially abundant (AGER and MFAP2), while HHIP and FAM13A were not detected in mass spectrometry proteomics of LTRC lung tissue. **(B**) The residual correlation matrix of the proteins in the core network as calculated in Results 6. The color gradient represents the Spearman r; squares -new experimental interactions; circles -HUBRIS interactions; for more details see Figure 6.

This core network consists of eight proteins, including five COPD GWAS gene products (AGER, FBXO38, HHIP, FAM13A and MFAP2) and three non-GWAS hubs (TGM2, CAVIN1, and HSPA1A) that together bridge five of the six GWAS proteins into a single connected module (Figure 6A). These three hubs are newly identified interactors that connect to two or more COPD GWAS gene products and exhibit significant differences in protein abundance between COPD and control lung tissue. Two GWAS gene products within the network, AGER and MFAP2, were differentially abundant in COPD lung tissue, with AGER showing lower and MFAP2 higher protein levels compared to controls. For HHIP and FAM13A, no proteomic signal could be measured. TGM2, identified in 16HBE cells, was increased in COPD lung tissue. In contrast, CAVIN1, identified in IMR90 cells, and HSPA1A, identified as an interactor in both 16HBE and IMR90 cells, exhibited reduced protein levels in COPD compared to controls. HSPA1A is also within 1 Mb of rs2070600, a COPD GWAS SNP and coding variant within *AGER*.

The residual correlation matrix of this core network (Figure 7B) confirms that several of these physical-interaction edges are accompanied by significant proteomic correlation. Despite its strong proteomic correlation, we did not find experimental evidence of AGER to be directly interacting with CAVIN1; as noted above, AGER does directly interact with another caveolar protein, EHD2. Moreover, the AGER-centered caveolar correlation block and the MFAP2 –TGM2 block from Figure 6 are conserved within this core network, suggesting strong functional connectivity.

### 9. Discussion

In this study, we combined affinity purification mass spectrometry (AP-MS), cell type-contextualized protein-protein interaction networks, and lung tissue transcriptomics and proteomics data to investigate how multiple COPD GWAS gene products collectively contribute to disease-relevant molecular networks. By experimentally mapping interactions for six well-established COPD GWAS genes in lung-relevant epithelial (16HBE) and fibroblast (IMR90) cell lines, we identified hundreds of previously unreported protein interactions and demonstrated that these interactions significantly impact our understanding of the network architecture underlying COPD risk.

The newly identified interactors and the subsequently generated cell type-specific subnetworks display stark cell type-specificity. These patterns emphasize the importance of tissue context in mapping the interaction landscape of COPD GWAS genes –and likely other complex diseases as well. Moreover, the number of newly identified interactors did not reflect the existing data in public PPI databases. This discrepancy highlights the complementary need for experimentally derived, cell type-specific interaction data and the limitations of relying on non-specific PPI networks for disease modeling. A potential caveat, however, is that AP-MS data has a substantial false-negative rate, so distinguishing cell type-specificity from incomplete coverage is quite difficult with the available data.

One of our key findings is that incorporating newly identified interactions significantly reduced the network distance among the six COPD GWAS gene products in the cell type-specific networks; however, this was observed only when the interactions were filtered based on the CRAPome database. We also observed that this reduction in network distance was not driven by random increases in connectivity but instead by a subset of proteins that interact with multiple GWAS gene products, effectively bridging distinct genetic risk loci. Pathway enrichment analysis of the multi-GWAS interactor network revealed both cell type-specific and shared biological processes previously implicated in COPD. The only protein that showed bridging properties in both cell types is THBS1, a gene that has also been identified in genetic association studies of lung function [35], functionally implicated in idiopathic pulmonary fibrosis (IPF) [36] and pulmonary hypertension [37].

By integrating several layers of multi-omic data and network properties, we refined an initial network of newly identified PPIs to a compact core module consisting of eight proteins that bridge multiple GWAS gene products and show differential protein abundance in COPD lungs. This core includes five GWAS genes (*AGER, FBXO38, FAM13A, HHIP,* and *MFAP2*) and three non-GWAS proteins: the caveolar protein CAVIN1, the ECM crosslinking enzyme TGM2, and the heat-shock protein HSPA1A. We address each of these in the following paragraphs.

Chen showed that *Cav1* knockout mice have increased susceptibility to develop emphysema and increased autophagy with chronic cigarette smoke (CS) exposure [38]. However, Volonte found that *Cav1* knockout mice had reduced sensitivity to develop emphysema with chronic CS exposure [39]. Thus, the directionality of caveolar perturbations on emphysema susceptibility is uncertain. AGER encodes sRAGE, a protein biomarker strongly associated with emphysema [40], and three independent layers of data from our work converge on the combined relevance of AGER and caveolae in COPD. First, we see striking **residual correlation** between the caveolar proteins CAVIN1, EHD1, EHD2, and CAV1 with AGER. Both RAGE [41,42] and caveolar proteins [43] are type 1 alveolar epithelial (AT1) markers that are highly expressed in AT1 but not in type-2 alveolar epithelial (AT2) cells [44]. This indicates shared regulatory programs [45] as well as potential spatial co-localization, which has already been suggested in endothelial cells [46]. Second, the above mechanisms are also supported by the direct **protein-protein interaction** we identified between AGER and EHD2 in IMR90 cells. Finally, these same caveolar proteins were among the most significantly **differentially abundant** in COPD lungs (Madha-Krause, in preparation). The lower levels of caveolar (and AGER) protein abundance in COPD may be an indicator of AT1 depletion in emphysema [47], but it is also possible that by relaying the genetic perturbation of GWAS genes, caveolae may play a more direct role in disease pathogenesis. In our previous study, we confirmed the interaction between CAVIN1 and HHIP via co-immunoprecipitation [29], while in this study we identified the CAVIN1 –MFAP2 interaction, which links caveolar proteins to several GWAS gene products. Overall, the caveolar signal convergence across interaction, differential abundance, and correlation marks a stark feature that may be key to understanding COPD pathobiology.

TGM2 is a hub that interacts with MFAP2 and FBXO38 in our lung-specific experimental data (also with AGER in the SAINTexpress-only, no CRAPome filtering scheme); however, in the HUBRIS reference database we found that it also connects to the other two non-GWAS hubs in the core network: CAVIN1 and HSPA1A. It is also the only newly identified interactor that has a significant **differential co-abundance** with one of the GWAS genes, shifting from a positive correlation with AGER in controls to a negative correlation in COPD. TGM2 is also one of the **top differentially abundant** proteins in our proteomic data, exhibiting significantly higher levels in COPD lungs. Transglutaminase 2 (encoded by *TGM2*) is an enzyme that crosslinks ECM proteins such as collagen and elastin to increase ECM stability [48], and its inhibition has been suggested as potential therapy for idiopathic pulmonary fibrosis (IPF) [49,50]. Elevated TGM2 in COPD lung tissue has been previously reported by Ohlmeier and colleagues [51]. TGM2 also promotes the epithelial-mesenchymal transition (EMT) [52,53], which we showed in our recent work may be a mechanism of aberrant wound healing in COPD [54]. TGM2 may be at the intersection bridging multiple COPD risk modules (MFAP2, AGER, caveolae), and its differential behavior makes it a prime candidate for future functional and mechanistic models of COPD.

HSPA1A (member of the HSP70 family) is the **most connected** of the three bridging nodes; however, its expression does not strongly correlate with any of the GWAS gene products, which reinforces its canonical role as a more generic stress-response protein. HSPA1A has been implicated in COPD by several studies. Xie et al. showed that low levels of Hsp70 in airway smooth muscle were associated with COPD severity [55]. In contrast, Dong et al. found that HSP70 expression was *increased* in the lung tissues of COPD patients and current smokers, and cigarette smoke induced the mRNA and protein expression of HSP70 in 16HBE cells [56]. Hlapčić et al. also found a positive correlation between HSP70 levels and COPD severity, and suggested that HSP70 may modulate immune response through RAGE signaling [57]. This latter result is supported by the HSPA1A -AGER PPI we identified; however, our proteomic data show *lower* levels of HSPA1A in COPD lungs. The apparent inconsistency within the literature and with our own results suggests that the role of HSPA1A may be highly context-specific.

Together, the above-discussed hubs define a compact core network that connects five of the six COPD GWAS proteins through a small number of experimentally identified, disease relevant interactions. Co-abundance data suggest that this network connects two key COPD-relevant functional modules: a caveolar block linked to AGER and an ECM/tissue remodeling block linked to MFAP2, with TGM2 being a potential disease-sensitive switch. All these programs are essential for alveolar differentiation and tissue repair, which is a clear target of current COPD research. Future investigation should involve determining the directionality, biochemical, and functional nature of these interactions and the effect of genetic perturbations (GWAS risk) on their canonical function.

Several limitations should be considered when interpreting our findings. The AP-MS experiments were performed in immortalized cell lines, which do not fully recapitulate *in vivo* lung biology. However, the use of two lung-relevant cell types and integration with human lung proteomic data may mitigate this concern. Due to the large number of interactions and our previous success in validating interactions with co-immunoprecipitation [29], we did not perform co-immunoprecipitation validation in this study. Although we used rigorous experimental and analytical approaches to detect protein-protein interactions, additional (potentially weaker) interactions were likely missed by our approach, which also limits our cell type-specific claims. Additionally, while network proximity and differential abundance highlight candidate disease modules, functional perturbation studies will be required to identify the causal roles of individual interactions and proteins within COPD pathogenesis.

A further limitation is that only three of the six COPD GWAS proteins (AGER, MFAP2, FBXO38) were detected in the lung proteomic data, so the correlation and differential abundance evidence for the remaining risk gene products, HHIP, IREB2, and FAM13A, could not be assessed at the protein level. Moreover, the proteomic measurements were derived from bulk lung tissue samples; therefore, the cell type-specificity of the signal cannot be determined. Future research will be needed to determine whether different network relationships influence the heterogeneous disease manifestations of COPD.

Despite these limitations, this study demonstrates the value of protein-protein interaction assessment and a meaningful way to combine multiple layers of omics data. Our final core module can be a useful scaffold for generating hypotheses and motivating targeted perturbation experiments to understand biological mechanisms for COPD pathogenesis.

## Methods

### 1. Cell culture and AP-MS

Human bronchial epithelial 16HBE and embryonic kidney epithelial 293T cells were purchased from ATCC (Minnesota, VA). Human telomerase reverse transcriptase (hTERT)-immortalized human fetal lung IMR-90 cells were obtained from Dr William Hahn’s laboratory (Dana-Farber Cancer Institute). 16HBE and IMR90 cells were cultured in Eagle’s minimum essential medium (EMEM). 293T cells were maintained in Dulbecco’s modified Eagle’s medium (DMEM). EMEM and DMEM were supplemented with 10% fetal bovine serum (FBS), 100 U/mL penicillin, and 100 μg/mL streptomycin. Cells were maintained at 37 °C in a humidified incubator with 5% CO_2_.

For each of the six bait proteins, triplicate affinity purifications were performed using independently cultured cells infected with lentiviral constructs expressing hemagglutinin (HA)-tagged versions of the corresponding bait proteins or an empty vector control. Following blasticidin selection (5 µg/mL), protein complexes were affinity-purified from 2 mL of cell lysates (protein concentration of 8–10 mg/mL) using anti-HA agarose (A2095, Sigma-Aldrich) and analyzed by LC–MS/MS.

### 2. Evaluation of PPI interactions

Raw spectral count data were processed using SAINTexpress [32] to estimate the probability of bait-prey interactions. To further control for nonspecific and/or highly promiscuous interactors, we incorporated contaminant counts from the CRAPome database [33]. The CRAPome data was used as an additional filter to exclude proteins commonly detected across unrelated AP-MS experiments, thus identifying a more conservative set of interactions.

### 3. The HUBRIS network

The HUBRIS reference network combines 8 public PPI databases: HIPPIE [58], HumanNet [59], HURI [60], BioGrid [61], Reactome [62], Interactome3D [63], NCBI [64], and StringDB [65] (in StringDB, we only included edges with a non-zero experimental score). The merging procedure is described in detail in the Methods section of [29]. The largest connected component of the network contains N=20,103 nodes and E=896,648 edges. For all databases the date of download is 2025 January 6.

### 4. Cell type-specific filtering of HUBRIS based on RNA-Seq data

We utilized RNA-Seq data obtained from the two cell types we used in this study: IMR90 and 16HBE. The RNA-Seq data are accessible at [66].

For both cell lines, we used the same criteria: we used a cutoff of 0.5 on the log2 of the TPM values to identify expressed genes. The value of 0.5 was used to obtain an appropriate balance between removing nodes with low expression while still retaining all proteins measured in the AP-MS experiments.

Finally, to create the cell type-specific reference networks, we kept genes that were expressed and removed every other node from HUBRIS. We then identified the remaining largest connected component (LCC) to create the cell type-specific network. The 16HBE-specific network had N=12,603, E=657,356; the IMR90-specific network N=12,213, E=637,128 nodes and edges, respectively.

### 5. Network distance analysis

To assess whether the newly identified GWAS protein interactors decrease the network distance, we compared the mean pairwise shortest path length between the six GWAS gene products in the empirical network versus a degree-preserving null model.

#### Network construction

The empirical networks were constructed as a combination of the cell type-specific HUBRIS networks (filtered based on IMR90 or 16HBE RNA-Seq data, see Methods 3 and 4) and the newly identified edges from their respective cell lines. We generated separate networks with edges validated in SAINTexpress alone, and with SAINTexpress plus CRAPome filtering (see Methods 2). The resulting 4 networks were undirected and unweighted.

#### Distance metric

For each network we computed the shortest path length between all 15 unique pairs of the 6 GWAS gene products, and calculated the mean as the empirical mean distance (marked with *d* in the statistical analysis).

#### Null model

Adding edges to a network can only reduce the shortest path distances, so we needed a null model to assess the expected distance reduction. We did this by adding an equal number of edges connected to the same 6 GWAS gene products on one end (we were deliberately sampling interactors of these proteins) and nodes of similar degree on the other.

We generated *n_perm_* = 1000 networks where we rewired the non-GWAS endpoint of every newly added experimental edge, and connected it to a node drawn from the same degree bin as the empirical endpoint node. The node degree bins were defined as deciles of the HUBRIS network degree distribution. This way, the GWAS endpoints, the total number of edges, all the HUBRIS edges (baseline topology) and the assortativity of the new edges was held fixed.

#### Significance

For each cell line and filtering criterion, we computed a standard Z score: *Z* = (*d_emp_*− < *d_null_*>)/σ*_null_*, where *d_emp_* is the empirical mean distance between the GWAS gene pairs, < *d_null_*> is the average mean distance across all null simulations, and σ*_null_* is the standard deviation of *d_null_* across all null simulations.

The one-sided empirical p-value was calculated as the proportion of randomizations whose mean shortest-path length was less than or equal to the empirical value:

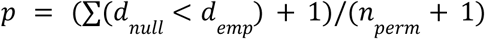

### 6. Proteomic data and covariate adjustment

Bulk lung tissue proteomic data was generated from the Lung Tissue Research Consortium (LTRC) (Madha-Krause, in preparation), across 4,511 proteins from 494 COPD and 364 control lung tissue samples. Of the six COPD GWAS gene products studied, only three -AGER, MFAP2, and FBXO38 -were detected at the protein level; therefore, all proteomic analyses were restricted to these three GWAS proteins (out of 6).

The differential abundance analysis was conducted using the limma package [67], controlling for age, sex, smoking status, smoking pack-years, ten ancestry principal components, and fifteen surrogate variables (Madha-Krause, in preparation). Adjusted p-values below 0.05 were considered significant.

### 7. Residual correlation and differential correlation analysis

In the residual correlation analysis, the ordinary least squares (OLS) method was used to fit the covariates (age, sex, smoking status, smoking pack-years, ten ancestry principal components, and fifteen surrogate variables). For each protein *i* the abundance *y_i_* across all samples was estimated as a linear function of the covariate matrix *X*:

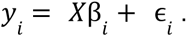

The residuals were calculated by subtracting the fitted from the observed values. All correlation (co-abundance) analyses were performed using the residual abundance values. Across the full proteome, the covariate set explained a median of 44.7% of per-protein variance; for the three GWAS proteins, the percent variance explained was: AGER 31%, MFAP2 27%, FBXO38 39%.

For each of the three measured GWAS proteins (AGER, MFAP2, FBXO38), we computed Spearman rank correlations between their covariate-adjusted residuals and the residuals of the other 4,505 proteins. P-values were adjusted for multiple testing for all three GWAS proteins using the Benjamini-Hochberg method across all 4,511 tested proteins. Adjusted p-values below 0.05 were considered significant.

For the differential correlation analysis, we were interested in how the COPD GWAS protein association depended on disease status. We calculated the Spearman r correlation for each GWAS protein pair separately across the COPD and control groups. We used the Fisher z-transformation, *z* = ½ *ln*[(1 + *r*) / (1 − *r*)], on the coefficients to approximate a normal-like distribution, and the difference between groups was calculated with the test statistic:

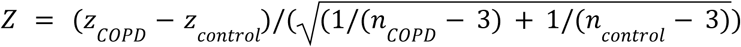

Two-sided p-values were calculated based on Z and adjusted for multiple testing with the Benjamini-Hochberg method.

### 8. PPI -correlation enrichment

To test whether the newly identified experimental interactors are enriched among the GWAS-protein correlates, we used the absolute value of the residual Spearman r to rank the proteins for each GWAS protein (AGER, MFAP2, FBXO38) in each cell line (IMR90, 16HBE). We then compared the rank distribution of the newly identified (CRAPome filtered) PPIs from each group with the non-interacting proteins using a one-sided Wilcoxon rank-sum test, under the alternative hypothesis that interactors rank higher (i.e., are more strongly correlated) than expected by chance.

### 9. Pathway enrichment analysis

The enrichment of the multi-GWAS bridging protein sets (Figure 4) was assessed by hypergeometric test (overrepresentation) against the cell type-specific background (see Methods 4), using the MSigDB Hallmark 2020 and KEGG 2021 Human libraries, with Benjamini-Hochberg correction using the GSEApy Python library.

For the correlation-enrichment (Figure 5) all measured proteins were ranked by their signed residual Spearman r and evaluated using preranked GSEA (GSEApy; 10,000 permutations, gene-set sizes 15–500) against MSigDB Hallmark 2020, GO Cellular Component 2025, and GO Biological Process 2026. Terms with BH adjusted p < 0.05 were significant. For visualization only, adjusted p-values were clipped at 1e-4, the permutation resolution limit.

### 10. Software versions

Python 3.11.15; pandas 3.0.1; NumPy 2.4.6; SciPy 1.17.1; statsmodels 0.14.6; scikit-learn 1.8.0; GSEApy 1.1.12; NetworkX 3.6.1

---

**Figure S1.**
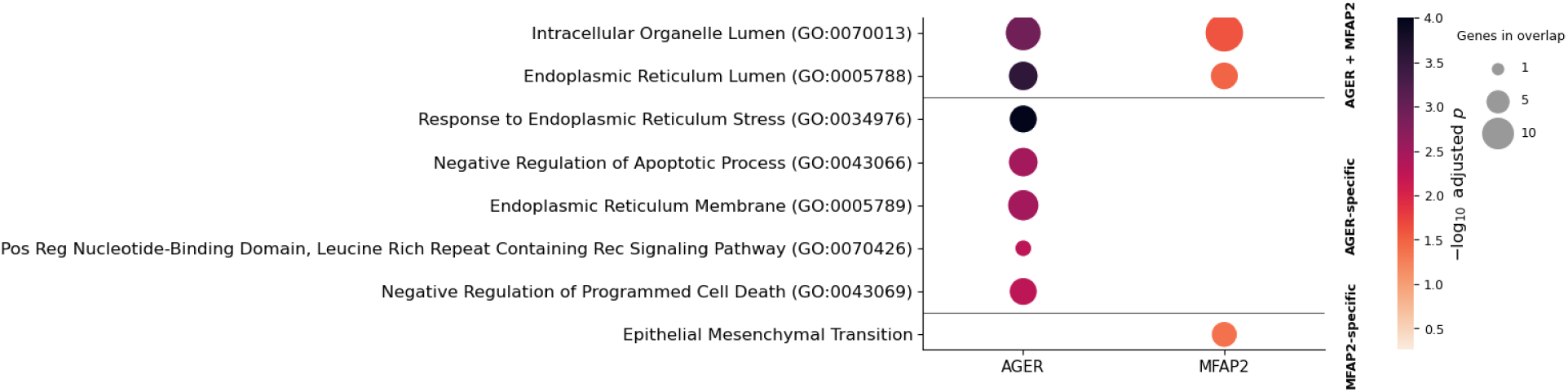
Pathway enrichment of new interactors that are also significantly correlated with AGER or MFAP2. Gene-set enrichment analysis (GSEA preranked, 10,000 permutations) of each GWAS protein’s genome-wide residual correlations, ranked by signed Spearman correlation coefficient, against MSigDB Hallmark 2020, GO Cellular Component 2025, and GO Biological Process 2026. Enriched terms (FDR < 0.05) are grouped into blocks by the combination of GWAS proteins in which they are significant with the top five terms shown per block, ranked within each block by the minimum −log₁₀(FDR) across the contributing GWAS proteins. Dot color indicates −log₁₀(FDR-adjusted p), capped at the permutation resolution limit (FDR floor = 1e-04), and dot size indicates the number of leading-edge genes. Only terms with FDR < 0.05 are displayed.

## References

1. Vos T. Global burden of chronic obstructive pulmonary disease. Lancet. 2020.

2. Adeloye D, Song P, Zhu Y, Campbell H, Sheikh A, Rudan I, et al. Global, regional, and national prevalence of, and risk factors for, chronic obstructive pulmonary disease (COPD) in 2019: a systematic review and modelling analysis. Lancet Respir Med. 2022;10: 447–458.

3. Agustí A, Hogg JC. Update on the pathogenesis of chronic obstructive pulmonary disease. N Engl J Med. 2019;381: 1248–1256.

4. Silverman EK. Genetics of COPD. Annu Rev Physiol. 2020;82: 413–431.

5. Hobbs BD, de Jong K, Lamontagne M, Bossé Y, Shrine N, Artigas MS, et al. Genetic loci associated with chronic obstructive pulmonary disease overlap with loci for lung function and pulmonary fibrosis. Nat Genet. 2017;49: 426–432.

6. Pillai SG, Ge D, Zhu G, Kong X, Shianna KV, Need AC, et al. A genome-wide association study in chronic obstructive pulmonary disease (COPD): identification of two major susceptibility loci. PLoS Genet. 2009;5: e1000421.

7. Cho MH, McDonald M-LN, Zhou X, Mattheisen M, Castaldi PJ, Hersh CP, et al. Risk loci for chronic obstructive pulmonary disease: a genome-wide association study and meta-analysis. Lancet Respir Med. 2014;2: 214–225.

8. Cho MH, Castaldi PJ, Hersh CP, Hobbs BD, Barr RG, Tal-Singer R, et al. A genome-wide association study of emphysema and airway quantitative imaging phenotypes. Am J Respir Crit Care Med. 2015;192: 559–569.

9. Zhou X, Baron RM, Hardin M, Cho MH, Zielinski J, Hawrylkiewicz I, et al. Identification of a chronic obstructive pulmonary disease genetic determinant that regulates HHIP. Hum Mol Genet. 2012;21: 1325–1335.

10. Castaldi PJ, Guo F, Qiao D, Du F, Naing ZZC, Li Y, et al. Identification of functional variants in the FAM13A chronic obstructive pulmonary disease genome-wide association study locus by massively parallel reporter assays. Am J Respir Crit Care Med. 2019;199: 52–61.

11. Cloonan SM, Glass K, Laucho-Contreras ME, Bhashyam AR, Cervo M, Pabón MA, et al. Mitochondrial iron chelation ameliorates cigarette smoke-induced bronchitis and emphysema in mice. Nat Med. 2016;22: 163–174.

12. Sambamurthy N, Leme AS, Oury TD, Shapiro SD. The receptor for advanced glycation end products (RAGE) contributes to the progression of emphysema in mice. PLoS One. 2015;10: e0118979.

13. Mecham RP, Gibson MA. The microfibril-associated glycoproteins (MAGPs) and the microfibrillar niche. Matrix Biol. 2015;47: 13–33.

14. Saferali A, Yun JH, Parker MM, Sakornsakolpat P, Chase RP, Lamb A, et al. Analysis of genetically driven alternative splicing identifies FBXO38 as a novel COPD susceptibility gene. PLoS Genet. 2019;15: e1008229.

15. Werder RB, Liu T, Abo KM, Lindstrom-Vautrin J, Villacorta-Martin C, Huang J, et al. CRISPR interference interrogation of COPD GWAS genes reveals the functional significance of desmoplakin in iPSC-derived alveolar epithelial cells. Sci Adv. 2022;8: eabo6566.

16. Nakamura T, Lozano PR, Ikeda Y, Iwanaga Y, Hinek A, Minamisawa S, et al. Fibulin-5/DANCE is essential for elastogenesis in vivo. Nature. 2002;415: 171–175.

17. Wert SE, Yoshida M, LeVine AM, Ikegami M, Jones T, Ross GF, et al. Increased metalloproteinase activity, oxidant production, and emphysema in surfactant protein D gene-inactivated mice. Proc Natl Acad Sci U S A. 2000;97: 5972–5977.

18. Miller PG, Qiao D, Rojas-Quintero J, Honigberg MC, Sperling AS, Gibson CJ, et al. Association of clonal hematopoiesis with chronic obstructive pulmonary disease. Blood. 2022;139: 357–368.

19. Parker MM, Hao Y, Guo F, Pham B, Chase R, Platig J, et al. Identification of an emphysema-associated genetic variant near TGFB2 with regulatory effects in lung fibroblasts. Elife. 2019;8. doi:10.7554/eLife.42720

20. Hautamaki RD, Kobayashi DK, Senior RM, Shapiro SD. Requirement for macrophage elastase for cigarette smoke-induced emphysema in mice. Science. 1997;277: 2002–2004.

21. D’Armiento J, Dalal SS, Okada Y, Berg RA, Chada K. Collagenase expression in the lungs of transgenic mice causes pulmonary emphysema. Cell. 1992;71: 955–961.

22. Lao T, Jiang Z, Yun J, Qiu W, Guo F, Huang C, et al. Hhip haploinsufficiency sensitizes mice to age-related emphysema. Proc Natl Acad Sci U S A. 2016;113: E4681–7.

23. Yun JH, Lee C, Liu T, Liu S, Kim EY, Xu S, et al. Hedgehog interacting protein-expressing lung fibroblasts suppress lymphocytic inflammation in mice. JCI Insight. 2021;6. doi:10.1172/jci.insight.144575

24. Barabási A-L, Gulbahce N, Loscalzo J. Network medicine: a network-based approach to human disease. Nat Rev Genet. 2011;12: 56–68.

25. Menche J, Sharma A, Kitsak M, Ghiassian SD, Vidal M, Loscalzo J, et al. Disease networks. Uncovering disease-disease relationships through the incomplete interactome. Science. 2015;347: 1257601.

26. Pintacuda G, Hsu Y-HH, Tsafou K, Li KW, Martín JM, Riseman J, et al. Protein interaction studies in human induced neurons indicate convergent biology underlying autism spectrum disorders. Cell Genom. 2023;3: 100250.

27. Hsu Y-HH, Pintacuda G, Liu R, Nacu E, Kim A, Tsafou K, et al. Using brain cell-type-specific protein interactomes to interpret neurodevelopmental genetic signals in schizophrenia. iScience. 2023;26: 106701.

28. Morrow JD, Zhou X, Lao T, Jiang Z, DeMeo DL, Cho MH, et al. Functional interactors of three genome-wide association study genes are differentially expressed in severe chronic obstructive pulmonary disease lung tissue. Sci Rep. 2017;7: 44232.

29. Deritei D, Inuzuka H, Castaldi PJ, Yun JH, Xu Z, Anamika WJ, et al. HHIP protein interactions in lung cells provide insight into COPD pathogenesis. Hum Mol Genet. 2025;34: 777–789.

30. Deritei D. HUBRIS -protein-protein interaction (PPI) networks. Zenodo; 2025. doi:10.5281/ZENODO.14604608

31. Lung Tissue Research Consortium (LTRC). In: NHLBI, NIH [Internet]. [cited 26 Jan 2026]. Available: https://www.nhlbi.nih.gov/science/lung-tissue-research-consortium-ltrc

32. Teo G, Liu G, Zhang J, Nesvizhskii AI, Gingras A-C, Choi H. SAINTexpress: improvements and additional features in Significance Analysis of INTeractome software. J Proteomics. 2014;100: 37–43.

33. Mellacheruvu D, Wright Z, Couzens AL, Lambert J-P, St-Denis NA, Li T, et al. The CRAPome: a contaminant repository for affinity purification-mass spectrometry data. Nat Methods. 2013;10: 730–736.

34. Sakornsakolpat P, Prokopenko D, Lamontagne M, Reeve NF, Guyatt AL, Jackson VE, et al. Genetic landscape of chronic obstructive pulmonary disease identifies heterogeneous cell-type and phenotype associations. Nat Genet. 2019;51: 494–505.

35. Shrine N, Izquierdo AG, Chen J, Packer R, Hall RJ, Guyatt AL, et al. Multi-ancestry genome-wide association analyses improve resolution of genes and pathways influencing lung function and chronic obstructive pulmonary disease risk. Nat Genet. 2023;55: 410–422.

36. Walsh SM, Worrell JC, Fabre A, Hinz B, Kane R, Keane MP. Novel differences in gene expression and functional capabilities of myofibroblast populations in idiopathic pulmonary fibrosis. Am J Physiol Lung Cell Mol Physiol. 2018;315: L697–L710.

37. Peng B, Zhou Y, Fu X, Chen L, Pan Z, Yi Q, et al. THBS1 mediates hypoxia driven EndMT in pulmonary hypertension. Pulm Circ. 2024;14: e70019.

38. Chen Z-H, Lam HC, Jin Y, Kim H-P, Cao J, Lee S-J, et al. Autophagy protein microtubule-associated protein 1 light chain-3B (LC3B) activates extrinsic apoptosis during cigarette smoke-induced emphysema. Proc Natl Acad Sci U S A. 2010;107: 18880–18885.

39. Volonte D, Kahkonen B, Shapiro S, Di Y, Galbiati F. Caveolin-1 expression is required for the development of pulmonary emphysema through activation of the ATM-p53-p21 pathway. J Biol Chem. 2009;284: 5462–5466.

40. Yonchuk JG, Silverman EK, Bowler RP, Agustí A, Lomas DA, Miller BE, et al. Circulating soluble receptor for advanced glycation end products (sRAGE) as a biomarker of emphysema and the RAGE axis in the lung. Am J Respir Crit Care Med. 2015;192: 785–792.

41. Demling N, Ehrhardt C, Kasper M, Laue M, Knels L, Rieber EP. Promotion of cell adherence and spreading: a novel function of RAGE, the highly selective differentiation marker of human alveolar epithelial type I cells. Cell Tissue Res. 2006;323: 475–488.

42. Buckley ST, Ehrhardt C. The receptor for advanced glycation end products (RAGE) and the lung. J Biomed Biotechnol. 2010;2010: 917108.

43. Newman GR, Campbell L, von Ruhland C, Jasani B, Gumbleton M. Caveolin and its cellular and subcellular immunolocalisation in lung alveolar epithelium: implications for alveolar epithelial type I cell function. Cell Tissue Res. 1999;295: 111–120.

44. Jung K, Schlenz H, Krasteva G, Mühlfeld C. Alveolar epithelial type II cells and their microenvironment in the caveolin-1-deficient mouse. Anat Rec (Hoboken). 2012;295: 196–200.

45. Zhao W, Lin Y, Xiong J, Wang Y, Huang G, Deng Q, et al. RAGE mediates β-catenin stabilization via activation of the Src/p-Cav-1 axis in a chemical-induced asthma model. Toxicol Lett. 2018;299: 149–158.

46. Varshavskaya KB, Petrushanko IY, Mitkevich VA, Barykin EP, Makarov AA. Post-translational modifications of beta-amyloid alter its transport in the blood-brain barrier in vitro model. Front Mol Neurosci. 2024;17: 1362581.

47. Zhang Y, Wei H, Nouws J, Jiang W, Brewster RM, Nguyen JP, et al. Aberrant cellular communities underlying disease heterogeneity in chronic obstructive pulmonary disease. Nat Genet. 2026;58: 376–391.

48. Sanders YY, Liu G. Transglutaminase-2: Nature’s glue in lung fibrosis? Am J Respir Cell Mol Biol. 2021;65: 243–244.

49. Philp CJ, Siebeke I, Clements D, Miller S, Habgood A, John AE, et al. Extracellular matrix cross-linking enhances fibroblast growth and protects against matrix proteolysis in lung fibrosis. Am J Respir Cell Mol Biol. 2018;58: 594–603.

50. Olsen KC, Epa AP, Kulkarni AA, Kottmann RM, McCarthy CE, Johnson GV, et al. Inhibition of transglutaminase 2, a novel target for pulmonary fibrosis, by two small electrophilic molecules. Am J Respir Cell Mol Biol. 2014;50: 737–747.

51. Ohlmeier S, Nieminen P, Gao J, Kanerva T, Rönty M, Toljamo T, et al. Lung tissue proteomics identifies elevated transglutaminase 2 levels in stable chronic obstructive pulmonary disease. Am J Physiol Lung Cell Mol Physiol. 2016;310: L1155–65.

52. Kumar A, Xu J, Brady S, Gao H, Yu D, Reuben J, et al. Tissue transglutaminase promotes drug resistance and invasion by inducing mesenchymal transition in mammary epithelial cells. PLoS One. 2010;5: e13390.

53. Cao L, Shao M, Schilder J, Guise T, Mohammad KS, Matei D. Tissue transglutaminase links TGF-β, epithelial to mesenchymal transition and a stem cell phenotype in ovarian cancer. Oncogene. 2012;31: 2521–2534.

54. Deritei D, Anamika WJ, Zhou AX, Yun JH, Sauler M, Cho MH, et al. HHIP’s dynamic role in epithelial wound healing reveals a potential mechanism of COPD susceptibility. Proc Natl Acad Sci U S A. 2026;123: e2424377123.

55. Xie J, Zhao J, Xiao C, Xu Y, Yang S, Ni W. Reduced heat shock protein 70 in airway smooth muscle in patients with chronic obstructive pulmonary disease. Exp Lung Res. 2010;36: 219–226.

56. Dong J, Guo L, Liao Z, Zhang M, Zhang M, Wang T, et al. Increased expression of heat shock protein 70 in chronic obstructive pulmonary disease. Int Immunopharmacol. 2013;17: 885–893.

57. Hlapčić I, Hulina-Tomašković A, Grdić Rajković M, Popović-Grle S, Vukić Dugac A, Rumora L. Association of plasma heat shock protein 70 with disease severity, smoking and lung function of patients with chronic obstructive pulmonary disease. J Clin Med. 2020;9: E3097.

58. Alanis-Lobato G, Andrade-Navarro MA, Schaefer MH. HIPPIE v2.0: enhancing meaningfulness and reliability of protein-protein interaction networks. Nucleic Acids Res. 2017;45: D408–D414.

59. Kim CY, Baek S, Cha J, Yang S, Kim E, Marcotte EM, et al. HumanNet v3: an improved database of human gene networks for disease research. Nucleic Acids Res. 2022;50: D632–D639.

60. Luck K, Kim D-K, Lambourne L, Spirohn K, Begg BE, Bian W, et al. A reference map of the human binary protein interactome. Nature. 2020;580: 402–408.

61. Oughtred R, Rust J, Chang C, Breitkreutz B-J, Stark C, Willems A, et al. The BioGRID database: A comprehensive biomedical resource of curated protein, genetic, and chemical interactions. Protein Sci. 2021;30: 187–200.

62. Milacic M, Beavers D, Conley P, Gong C, Gillespie M, Griss J, et al. The reactome pathway knowledgebase 2024. Nucleic Acids Res. 2024;52: D672–D678.

63. Mosca R, Céol A, Aloy P. Interactome3D: adding structural details to protein networks. Nat Methods. 2013;10: 47–53.

64. Index of/gene/GeneRIF. [cited 14 July 2026]. Available: https://ftp.ncbi.nih.gov/gene/GeneRIF/.

65. Szklarczyk D, Kirsch R, Koutrouli M, Nastou K, Mehryary F, Hachilif R, et al. The STRING database in 2023: protein-protein association networks and functional enrichment analyses for any sequenced genome of interest. Nucleic Acids Res. 2023;51: D638–D646.

66. GEO Accession viewer. [cited 14 July 2026]. Available: https://www.ncbi.nlm.nih.gov/geo/query/acc.cgi?acc=GSE285360.

67. Ritchie ME, Phipson B, Wu D, Hu Y, Law CW, Shi W, et al. limma powers differential expression analyses for RNA-sequencing and microarray studies. Nucleic Acids Res. 2015;43: e47.

