## Supplementary figures and images for "Newly identified COPD GWAS protein interactors reveal a potential disease network module"

### Figure_S1.png

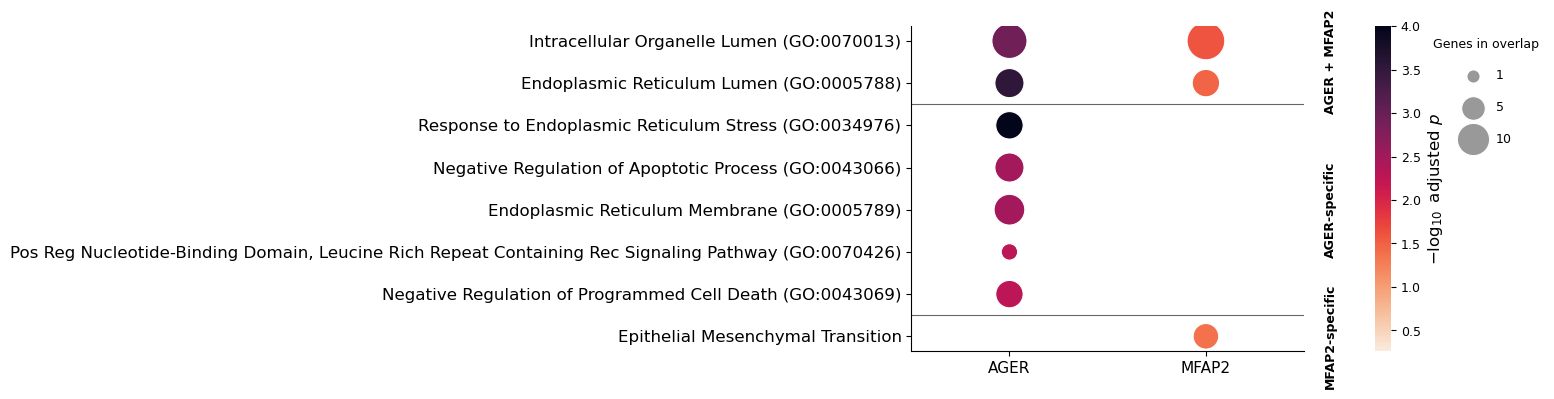
